# Experimental data supporting a novel hypothesis for the rhythmic initiation of proximal colon motor complexes

**DOI:** 10.1101/2024.12.24.630280

**Authors:** Wilmarie Morales-Soto, Emma S. Stiglitz, Brian S. Edwards, Kristen M. Smith-Edwards

**Author notes:** Corresponding author: Kristen M Smith-Edwards.

## Abstract

Current models of colon motility are largely based on studies of distal regions where distension-induced neural peristalsis predominates, but the proximal colon significantly differs in terms of cellular organization and its rhythmic motor activity that continues without external sensory input, emphasizing the need to define the unique mechanisms utilized by the proximal colon. With the long-term goal of developing a new model for the rhythmic initiation of proximal colon motor complexes (CMCs), we used *in situ* calcium imaging to define activity patterns in key players for colon motility [i.e., submucosal interstitial cells of Cajal (ICC-SM) and myenteric neurons of the enteric nervous system (ENS)], while simultaneously monitoring motor output in the proximal mouse colon. We observed repeated patterns of activity in ICC-SM and ENS myenteric neurons during the intervals between CMCs that could be used to predict the timing of subsequent CMC events. Based on our findings, we propose a novel hypothesis that cyclical interactions between the ENS and ICC-SM act as an intrinsic pattern generator for the rhythmic initiation of proximal CMCs.

## Introduction

The colon is tasked with handling, processing and moving luminal contents with physical properties that change along its length and, as such, is capability of generating diverse and region-specific motility behaviors. Proximal regions of the colon employ complex motor patterns that facilitate water reabsorption and the transformation of amorphous fecal material into solid and discrete pellets. Distal regions engage motor patterns that facilitate sustained propulsion of fecal pellets to the rectum for temporary storage and elimination. Colonic motor patterns result from smooth muscle contractions and relaxations driven by various patterns of activity in myenteric neurons of the enteric nervous system (ENS) and interstitial cells of Cajal (ICC; ‘pacemakers of the gut’). Each of these cellular components has been individually studied in the context of the neurogenic and myogenic motor patterns they generate, respectively.

However, colon motility is more likely a product of the integration of neuro-myogenic processes^1–3^, and how the ENS and ICC interact to coordinate the full complexity of colon motility behaviors is not well understood. Additionally, current models of colon motility are based on studies of distal regions^4, 5^, but the proximal colon has significant differences in terms of the ENS, ICC, and motor patterns^6–10^, emphasizing the need to define unique mechanisms utilized by proximal and distal regions of the colon that reflect their respective roles in fecal pellet *formation* versus *propulsion*.

Myogenic and neurogenic motor patterns have been measured in the colon using several animal models and in humans^1^. Ripples are myogenic contractions generated by slow waves of electrical activity in submucosal interstitial cells of Cajal (ICC-SM) that propagate via gap junctions into the circular smooth muscle^11^. The other major motor pattern, colon motor complexes (CMCs), consist of regular bouts of motor activity, require ENS activity (i.e., neurogenic), and appear as clusters of contractions that typically advance from proximal to distal colon^1^. Importantly, CMCs observed in proximal and distal regions exhibit several key differences^6^ and may represent two separate motor patterns driven by distinct mechanisms.

Existing models for the mechanisms underlying colon motility are based on distal regions, where neural peristalsis is initiated and self-sustained by a solid fecal pellet that distends the colon wall, sequentially activating the polarized ENS pathways responsible for propelling it forward^5, 12, 13^. In response to distension, intrinsic primary afferent neurons (IPANs) activate ascending excitatory and descending inhibitory neural networks to induce oral contraction and aboral relaxation of smooth muscle cells, resulting in forward movement of the pellet, distension and activation of the adjacent region. However, this “neuromechanical loop” model does not explain the regular rhythm of CMCs or how they are initiated in proximal regions where pellets have not yet formed^6^. Rather than distension-induced mechanisms, the spontaneous and regular generation of proximal CMCs indicates that this region of colon has an intrinsic ‘pacemaker’ capable of self-generating rhythmic CMCs, but the network mechanisms are unknown.

With the long-term goal of developing a new model for the rhythmic initiation of proximal CMCs, we used *in situ* calcium imaging to define activity patterns in ICC-SM and myenteric neurons of the ENS, while simultaneously monitoring ripples and CMCs in the proximal mouse colon. We observed repeated patterns of activity in ICC-SM and ENS myenteric neurons during the intervals between CMCs that could be used to predict the timing of subsequent CMC events. Following a CMC, a distinct group of myenteric neurons (Type 1) developed tonic spontaneous activity that continued for most of the inter-CMC interval. The activity across ICC-SM networks became more organized during this time, and activity in a different subset of neurons (Type 2), putative contraction-sensitive IPANs, increased as ICC-driven ripples became larger. Seconds prior to the next CMC, activity in Type 1 neurons ended and appeared to trigger robust activity in a third group of neurons (Type 3) and contractions of the CMC. Interestingly, distinct phases of ENS activity during CMC cycles corresponded to shifts in ICC frequency and organization across networks, and these dynamic changes in ICC were abolished when ENS activity was blocked. Although additional experiments are required to test our hypothesis, we propose a new model for the rhythmic initiation of proximal CMCs that involves cyclical, self-regenerating interactions among ICC-SM, a unique subset of IPANs that respond to ICC-driven ripples, and myenteric motor networks that feedback onto ICC-SM.

## Results

### CMCs occur regularly in proximal regions without luminal contents or external sensory input

To better understand the mechanisms underlying the rhythmic initiation of CMCs in proximal colon, we video-recorded motility in colon preparations that had been (1) emptied of luminal contents but remained closed or (2) cut open and pinned flat (**Fig 1A**). The second approach is more optimal for calcium imaging, and it ensures the complete absence of luminal stimulation by contents as well as pressure that can build up in the closed preparation. There were no differences in the frequency of CMC initiation (**Fig 1B**) or the length of propagation between closed and open preparations (**Fig 1C**). Stretching the colon tissue in open preparations did not significantly alter the length of propagation (**Fig 1D**) nor the frequency of CMCs in proximal colon (**Fig 1E**). However, the frequency of CMC events in distal colon was significantly higher in stretched colons (**Fig 1E**), supporting that the mechanism for CMCs in distal, but not proximal, colon is driven by distension-like sensory input. To further explore conditions that influenced CMCs, we also compared the frequency of CMC events in proximal versus distal colon when these regions were physically separated from one another by cutting the colon in half. In preparations with applied stretch, separating the proximal and distal colon had no effect on the frequency of CMCs (**Fig 1F**). However, in colon preparations that were not stretched, CMCs no longer occurred in distal colon when cut away from proximal colon, whereas CMCs continued to occur at similar frequencies in proximal colon without the distal colon intact (**Fig 1G**). Therefore, consistent with the concept that neural peristalsis drives motility in the distal colon, CMCs in distal regions seem to require stretch or distension of the colon wall, whereas the proximal colon appears to have an intrinsic pattern generator capable of producing regular CMCs without luminal input or distension.

**Figure 1.**
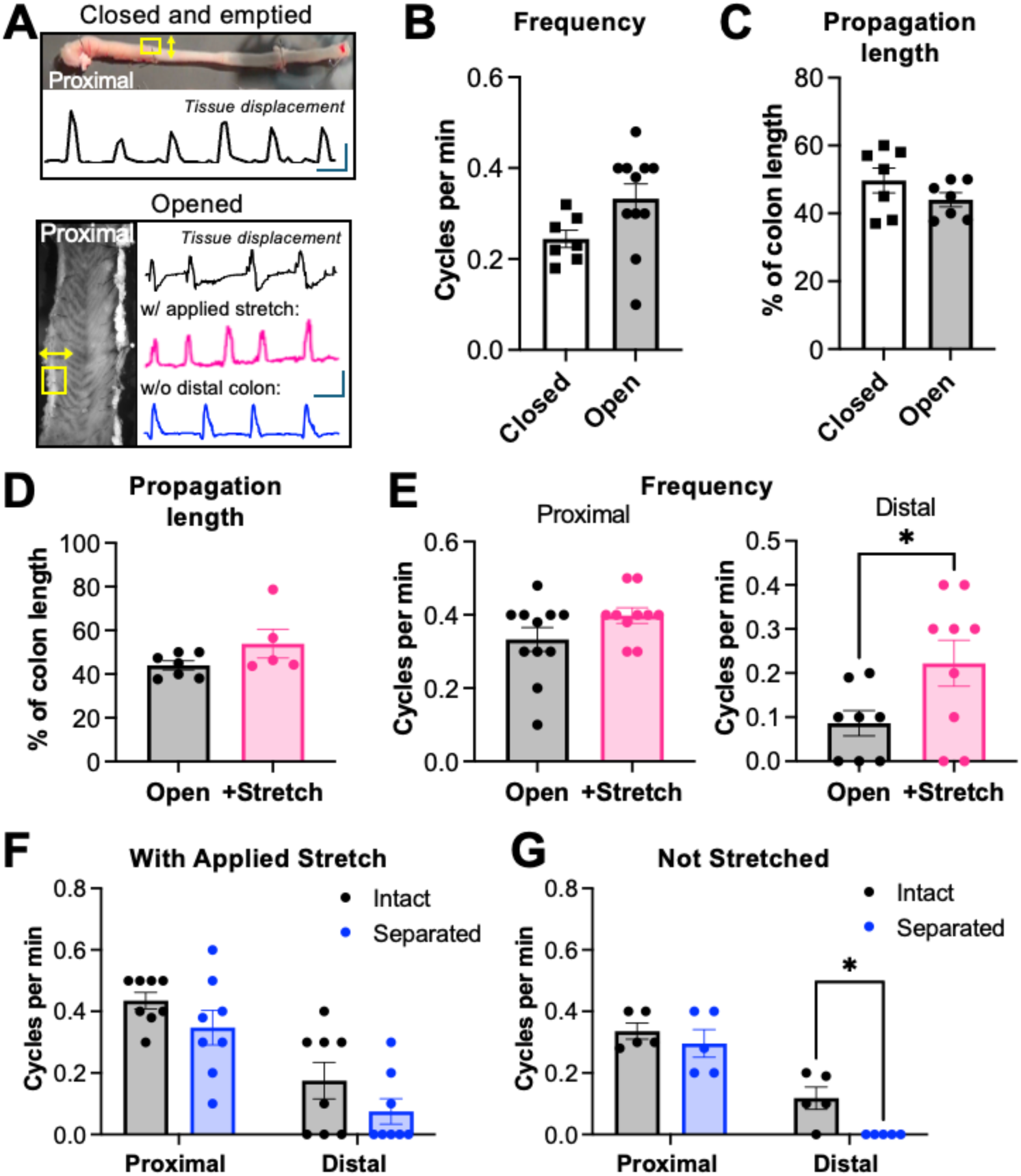
Proximal CMCs are rhythmic and do not require luminal contents. (A) Images of closed and open colon preparations and displacement traces of CMCs. (B) No changes in CMC frequency observed in closed and open colons (p>0.05, n=7-11, unpaired t-test). (C) No differences in the length of CMC propagation between closed and open colons (p>0.05, n=7, unpaired t-test). (D) No differences in length of propagation with applied stretch in the open colon preparation (p>0.05, n=5-7, unpaired t-test). (E) Stretching the open preparation increased CMC frequency in distal (p<0.05, n=8-9, unpaired t-test), but not proximal (p>0.05, n=10-11, unpaired t-test) colon. (F) When stretch was applied to the open preparation, CMC frequency remained similar in proximal and distal colon whether they were intact or separated (p>0.05, n=8, repeated measures two-way ANOVA). (G) Without applied stretch, CMC frequency decreased to zero when the distal colon was separated from the proximal half, whereas no changes were observed in the proximal colon (p<0.05, n=5, repeated measures two-way ANOVA).

### Dynamic changes in ICC-SM activity correlate to CMC initiation

In the colon, ICC-SM generate “pacemaker” slow waves of depolarization that drive smooth muscle ripple contractions^11^, but unlike myenteric ICC (ICC-MY) that are released from tonic nitrergic inhibition during CMCs^14, 15^, little is known about activity in ICC-SM networks relative to the timing of CMCs. To address this, we used Ca^2+^ imaging to measure ICC-SM network activity and ripples before, during and after CMCs initiated in proximal colon. The properties of ICC-SM activity underlying slow waves were visualized with timelapse color coded images, fluorescence intensity profiles of adjacent ROIs along the wave path, and spatiotemporal (ST) heat maps that were generated from time series recordings (**Fig 2A-B**).

**Figure 2.**
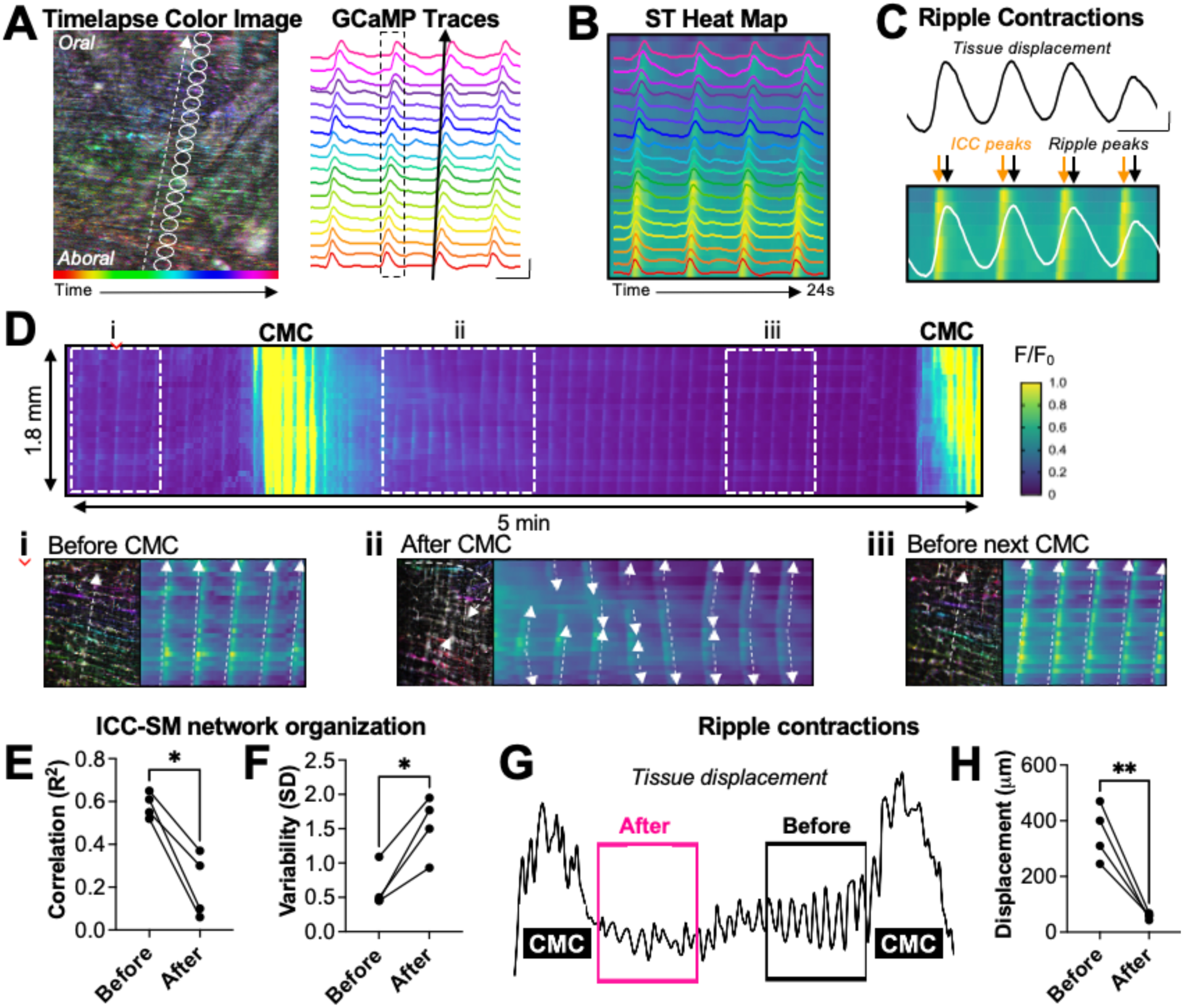
Repeated changes in ICC-SM network activity in proximal colon correspond to increased ripple contractions prior to CMCs. (A) Timelapse color coded image (ImageJ) showing adjacent ROIs drawn along the ICC-SM wave path traveling in the aboral to oral direction; GCaMP traces showing peaks in fluorescent signal from individual ROIs drawn over ICC-SM. (B) Example spatiotemporal heatmap using the intensity profiles from A. (C) By tracking tissue displacement, the ripple contractions generated by ICC-SM were also monitored. (D) Example ST heatmap of ICC-SM activity during two consecutive CMCs; i, organized waves before CMC; ii, disorganized waves after CMC; iii, reorganized waves before the next CMC. (E) Correlation coefficient (R^2) of GCaMP activity from adjacent ROIs was higher before versus after CMCs (p<0.05, paired t-test, n=4). (F) Variability (standard deviation) of intervals between ICC-SM oscillations was greater after versus before CMCs (p<0.05, paired t-test, n=4). (G) Example tissue displacement during and between two CMCs showed that (H) ripple contractions are greater before versus after CMCs (p<0.05, paired t-test, n=4).

The timelapse color code ImageJ plugin assigns a different color to each pixel in the image based on the timing of peak fluorescence, allowing an individual ICC-SM wave of activity to be visualized (**Fig 2A**). Adjacent ROIs were placed along the wave path, and the fluorescence intensity profiles were used to generate individual traces or spatiotemporal (ST) heat maps to view ICC-SM oscillations that repeatedly propagate along the wave path (**Fig 2B**). We also tracked the movement of the imaging field in the circular muscle (x) axis to measure the ripple contractions that followed each ICC-SM calcium wave (**Fig 2C**).

When imaging the same field of view to capture intervals between multiple CMCs, we observed dynamic and repeated changes in the organization of ICC-SM network activity relative to the timing of CMC events (**Fig 2D**). Before CMCs, activity in ICC-SM was well-organized, where Ca^2+^ waves propagated along the longitudinal axis from one end of the imaging field to the other for several cycles at regular intervals (**Fig 2Di**). Once a CMC passed the imaging field, ICC-SM activity was reduced and disorganized, where Ca^2+^ waves traveled opposite directions or crashed together, and the interval between cycles was not consistent (**Fig 2Dii**).

After some time, ICC-SM activity began to re-organize across the ICC-SM network, the next CMC soon followed (**Fig 2Diii**), and this pattern continued. In the 30 sec leading up to a CMC, the Ca^2+^ signals from adjacent ICC-SM displayed higher correlation (**Fig 2E**) and the intervals between ICC-SM Ca^2+^ waves were less variable (**Fig 2F**) compared to the 30 sec following a CMC. Further, the magnitude of the ripples generated by ICC-SM slow waves followed a similar pattern of cyclical changes relative to the timing of CMC events (**Fig 2G**). Before CMCs when ICC-SM network activity was most organized, the tissue displacement caused by ripple contractions was significantly greater than after CMCs, when ICC-SM network activity was disorganized (**Fig 2H**). Therefore, leading up to a CMC, ICC-SM activity becomes more and more organized, causing bigger and bigger ripples, but the CMC event itself (or cellular activity during the CMC) appears to act as a “brake” and resets the system by briefly disorganizing activity across ICC-SM networks.

### ENS activity influences ICC-SM networks in proximal colon

Previous studies have reported that ICC slow wave properties in the stomach and small intestine can be modulated by ENS activity via cholinergic and nitrergic signaling mechanisms^14–17^, and we hypothesized that the dynamic changes in colonic ICC-SM slow wave activity during CMC cycles are due to input from enteric neurons. To test this, we compared ICC-SM activity in the presence and absence of tetrodotoxin (TTX) to block ENS activity (Fig 3). Under baseline conditions, we observed similar changes described above in ICC-SM Ca^2+^ wave behavior during the intervals between CMCs (e.g., changes in direction of propagation, variability between cycles), but when ENS activity was blocked with TTX, ICC-SM Ca^2+^ waves displayed remarkable regularity across the field of view (**Fig 3B**). The basal frequency of ICC-SM Ca^2+^ waves in proximal colon regions significantly increased in the presence of TTX (**Fig 3C**), resulting in a decreased aboral-oral frequency gradient (i.e., smaller differences in frequency across the tissue) and more consistency in the direction of wave propagation compared to conditions with ongoing ENS activity. Further, the variability of intervals between ICC-SM Ca^2+^ waves was significantly lower in the presence of TTX (**Fig 3D**), indicating that ENS activity influences ICC-SM network activity in the proximal colon potentially by imposing frequency gradients.

**Figure 3.**
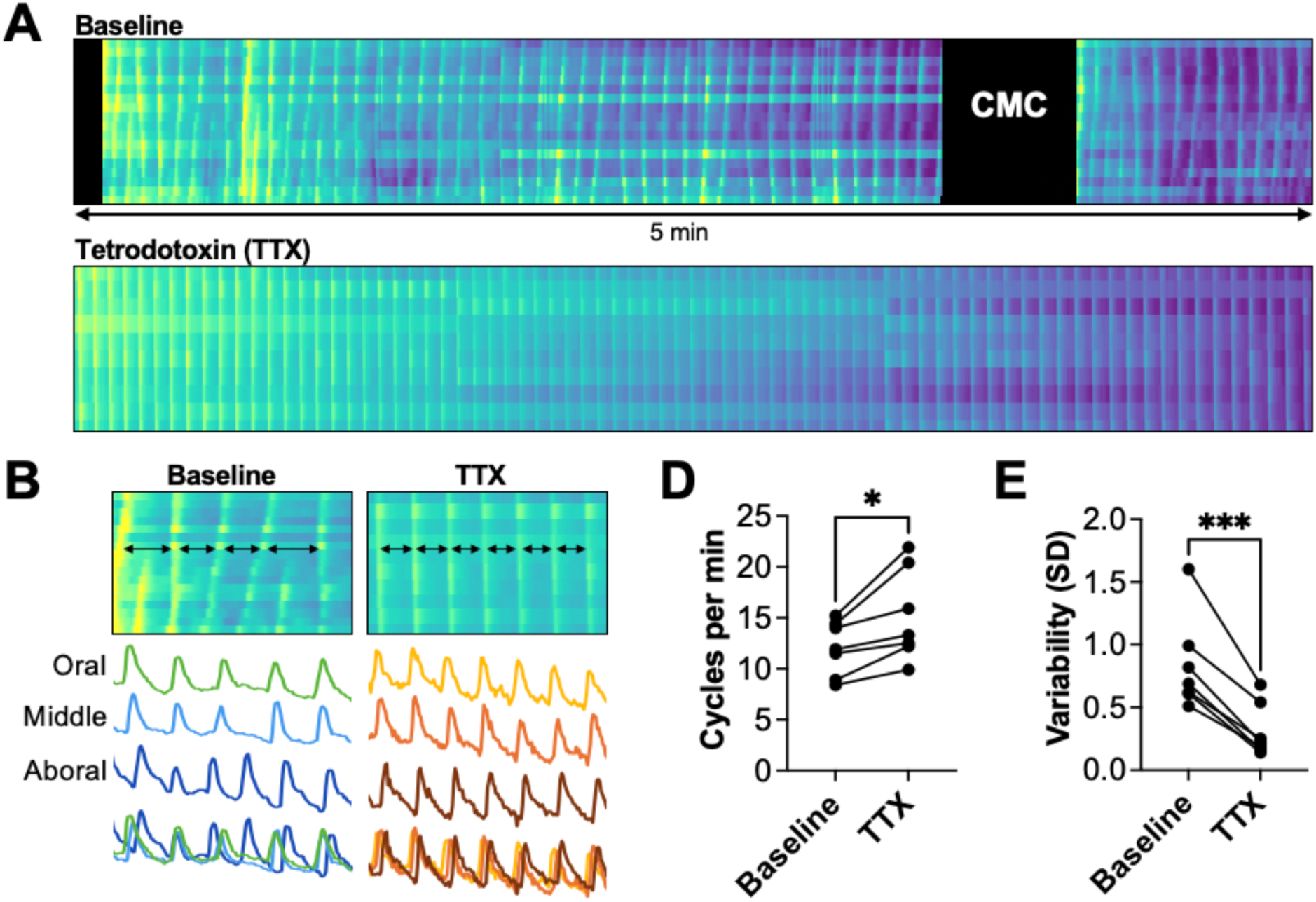
ICC-SM network activity is regular and lacks dynamic changes in absence of ENS activity. (A) Example spatiotemporal heatmaps of ICC-SM network activity during baseline conditions and when ENS activity is blocked with TTX. (B) Arrows show the irregular and regular spaced intervals between ICC-SM oscillations in baseline versus TTX conditions; these differences in organization are also observed in the GCaMP traces below. (D) ICC-SM frequency was increased (p<0.05, paired t-test, n=7) and (E) variability was decreased (<0.001, paired t-test, n=7) in TTX versus baseline.

### Three major patterns of activity in myenteric neurons of the ENS

To define patterns of ENS activity associated with rhythmic motility in the proximal colon, we next imaged activity in myenteric neurons before, during and after CMCs in the same field of view for 15-30 min. In the E2a-GCaMP model, the calcium indicator is genetically expressed in ICC and all neurons. Therefore, although the neurochemical phenotype of individual neurons could not be identified in real-time, we had the unique ability to relate the timing of activity patterns across all major cell types involved in CMCs. We observed three distinct patterns of activity relative to CMC timing (**Fig 4A**) that appeared to represent different functional classes of myenteric ENS neurons with unique roles for CMC initiation. The first group of neurons (‘Type 1) exhibited ongoing activity for most of the time between CMCs, but this activity turned off 5-10 sec before a CMC event. Another group of neurons (‘Type 3’) were active only during CMCs with little activity between events. Interestingly, the timing of activity in Type 1 and Type 3 neurons was non-overlapping; in other words, when Type 1 neurons were on, Type 3 neurons were off and vice versa, suggesting that inhibition of Type 1 neurons may be related to excitation of Type 3 neurons. Based on these characteristics, we predict that Type 1 neurons are nitrergic neurons that establish tonic inhibitory tone during the intervals between CMCs, whereas Type 3 neurons are motor neurons that must be engaged to produce contractions (and/or relaxations) during CMCs. The third group of neurons (‘Type 2’) displayed a pattern of activity with intermediary properties compared to the other groups. Type 2 neurons developed activity that increased in magnitude leading up to a CMC, and their activity temporally coincided with the increasing magnitude of ripple contractions evoked by ICC-SM, as if these neurons were specifically tuned to respond to myogenic contractions.

**Figure 4.**
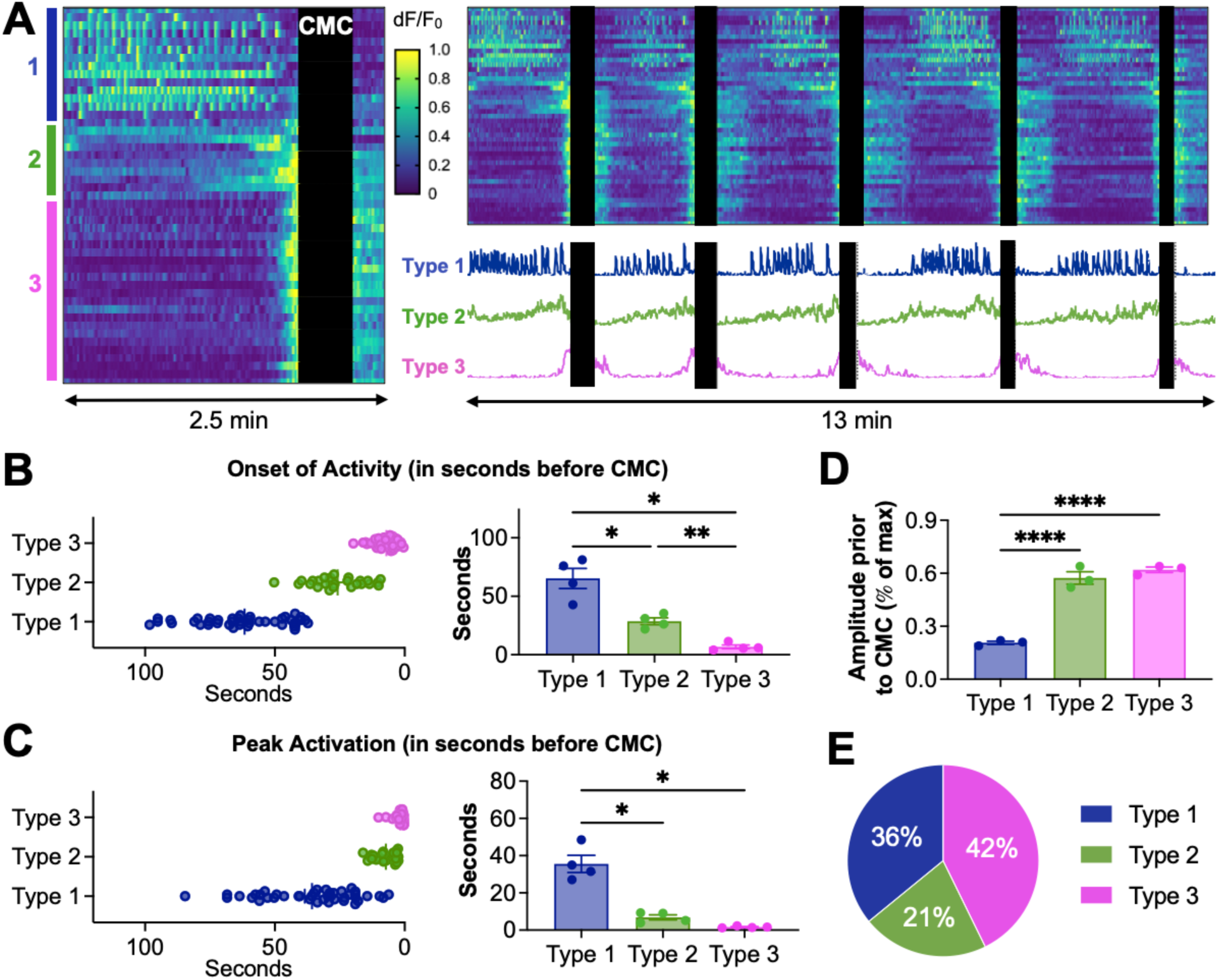
Three different activity patterns in myenteric neurons of the ENS during CMC cycles. (A) Each row in the ST heatmaps represents activity from individual neurons for 2.5 min (left) and 13 (right) min during CMC cycles (black boxes represent CMCs); as shown in the example traces, three different activity patterns that repeat each CMC cycle can be observed. (B) Relative to the next CMC, activity begins in Type 1 neurons earlier than the other types, and in Type 2 neurons earlier than Type 3, which are not activated until a few seconds before the CMC (p<0.05 and p<0.01, one-way ANOVA, n=4). (C) The peak activation of Type 2 and 3 neurons, occurring seconds before the CMC, was significantly different than Type 1 neurons (p<0.05, one-way ANOVA, n=4). (D) Amplitude of activity in Type 2 and 3 neurons before CMCs was significantly greater than in Type 1 neurons. (E) Proportions of neurons with Type 1, 2, and 3 activity.

We focused on three different characteristics to quantitatively categorize myenteric neurons based on the CMC activity patterns described above. The onset of activity relative to CMC timing differed significantly across the three subtypes and served as a useful defining feature to distinguish categories (**Fig 4B**). Peak activation in neurons with Type 2 and 3 activity patterns was within 10 seconds of the CMC and significantly lower compared to the timing of peak activation in neurons with Type 1 activity patterns (**Fig 4C**). In the seconds just before a CMC, activity in Type 1 neurons had significantly lower amplitude (∼20% of max) compared to Type 2 and 3 neurons (**Fig 4D**). Using these defining features, approximately 36% of myenteric neurons displayed Type 1 activity patterns, 21% Type 2, and 42% Type 3 (**Fig 4E**).

## Discussion

The regular occurrence of colon motor complexes or CMCs in several animal models has been appreciated for decades and commonly used to assess the functional output of the ENS. Thus, it is surprising that the mechanisms underlying the rhythmic and spontaneous initiation of CMCs have not yet been defined. Here, we used calcium imaging in full-thickness, full-length mouse colon to identify repeated activity patterns in the ENS and ICC-SM that are associated with CMCs in the proximal colon. Unlike the distal colon, where distension-evoked neural peristalsis predominates, the proximal colon appears to have an intrinsic ‘pacemaker’ capable of self-generating CMCs involving the cyclical interactions among ICC, IPANs, and the ENS.

### A novel hypothesis to explain proximal CMC rhythmicity

As shown in Fig 5, (1) ICC-SM slow waves organize across the proximal colon and ripple contractions get larger in magnitude. Once large enough, (2) ripples activate a subpopulation of contraction-sensitive intrinsic primary afferent neurons (IPANs) that (3) engage the ‘ENS program’ that is responsible for producing the CMC; Type 1 neurons turn off, Type 3 neurons turn on, and this switch in activity program is associated with the activation of ICC in the myenteric plexus (ICC-MY) and longitudinal muscle (LM). (4) Robust ENS activity during the CMC acts on ICC-SM, causing transient disorganization of their slow wave activity and disappearance of ripples. (5) This resets the cycle, where the next CMC depends on reorganization of ICC-SM, emergence of ripple contractions and subsequent activation of contraction-sensitive IPANs.

**Figure 5.**
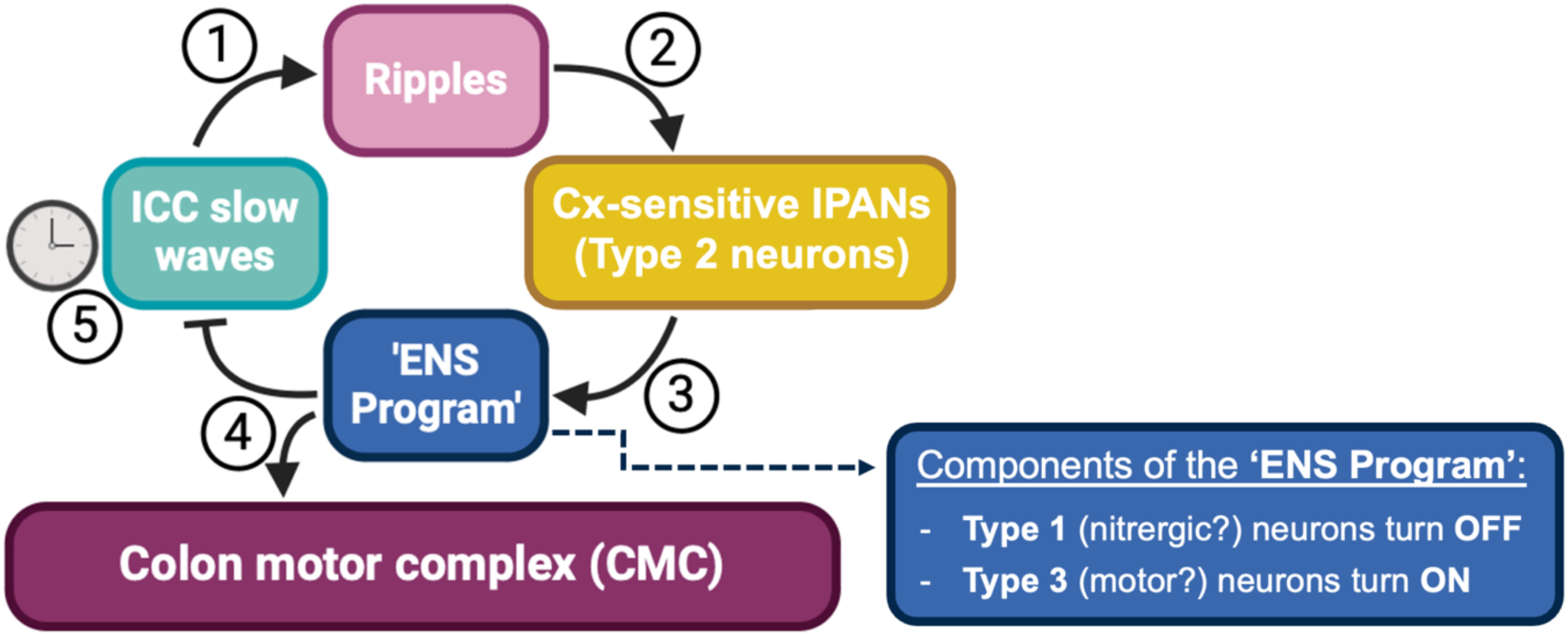
Proposed model for rhythmic CMCs in proximal colon. (1) ICC-SM calcium waves become organized, resulting in large ripples that (2) activate contraction-sensitive IPANs (Type 2 neurons); (3) IPANs engage the ‘ENS program’ that initiates a CMC (Type 1 neurons turn off and Type 3 neurons turn on). (4) Neural activity during the CMC disorganizes ICC-SM (5) which eventually re-organize over time, resetting the cycle.

Our proposed model is built upon two key principals that require further experimental and computational testing. For one, rather than the IPANs in distal colon that are activated in response to mechanical forces applied *extrinsically* (e.g., stretch or distension of the colon wall from a fecal pellet), we propose that a unique subset of IPANs, predominantly found in the proximal colon, respond to mechanical forces generated *intrinsically*, namely, the myogenic ripple contractions driven by ICC-SM. Once the molecular makeup of these Type 2 contraction-sensitive IPANs is identified, further functional characterization will be possible. For example, optogenetic studies can be used to determine how activation and/or inhibition alters CMC rhythmicity. Perhaps more important is testing whether these neurons are indeed contraction sensitive, but this will require approaches capable of evoking myogenic contractions on demand. The second key concept for our proposed model is neuronal feedback onto ICC-SM network activity. Previous studies in stomach, small intestine and colon have identified mechanisms by which cholinergic and nitrergic signaling alters ICC ^14–18^, but additional experiments are required to determine whether these mechanisms also apply to ICC-SM in the proximal colon.

Defining the cellular interactions that make up the intrinsic pattern generator for CMCs has been a challenge using traditional methods. Earlier studies relied on electrophysiology or chemical calcium dyes that required small, peeled pieces of tissue with minimal contractions and incomplete cellular networks. Genetically encoded calcium indicators allow the unique ability to visualize activity in individual neurons and non-neuronal excitable cells during ongoing contractions because the tissue does not require peeling and preparations can remain intact. As mentioned above, optogenetic approaches allow manipulation of activity in defined cell populations with superior spatial and temporal precision, thus eliminating off-target or compensatory effects that occur using pharmacology or genetic knock-out models. In future studies, optogenetics should be combined with calcium imaging to directly test the functional interactions between the ENS, ICC and IPANs that we propose contribute to CMC rhythmicity.

## Methods

### Animals

All work involving animals was conducted in accordance with the National Institutes of Health (NIH) Guide for Care and Use of Laboratory Animals and was approved by the Institutional Animal Care and Use Committee at the University of Pittsburgh and the Mayo Clinic (A00006603). Male and female mice with a C57BL/6 background at 10-16 weeks of age were used for all experiments. All mice were maintained on a 12-hour:12-hour light-dark cycle in a temperature-controlled environment with access to food and water ad libitum. Transgenic mice expressing the genetically encoded calcium indicator GCaMP6s in neuronal and non-neuronal cells (called E2aCre-GCaMP mice^10, 19, 20^) were bred in house and generated by crossing E2aCre mice (JAX#003724; RRID:IMSR_JAX:003724) with mice expressing a floxed-STOP-GCaMP6s sequence in the Rosa26 locus (JAX#028866; RRID:IMSR_JAX:028866).

### Measurement of Colon Motor Complexes (CMCs)

Mice were euthanized with isoflurane and full-length colons removed and placed into a Sylgard-lined dish (Dow Corning, Midland, MI) with circulating carboxygenated (95/5) artificial cerebral spinal fluid (ACSF) containing in mM: 117.9 NaCl, 4.7 KCl, 25 NaHCO3, 1.3 NaH2PO4, 1.2 MgSO47H2O, 2.5 CaCl2, 11.1 D-glucose, 2 sodium butyrate, and 20 sodium acetate (all purchased from Sigma-Aldrich, St. Louis, MO). For closed preparations, colons were pinned to the dish using minutien pins along the mesentery and gently flushed with ACSF to remove luminal contents. Intact closed colon preparations were video recorded using a Sony camcorder. For open preparations, the colons were cut open longitudinally, pinned flat using minutien pins with the mucosal side facing down, and motility was recorded on a Leica M205 FCA upright fluorescent dissecting microscope to obtain large fields of view (∼20mm). Several conditions were tested to determine the factors that influenced CMC rhythmicity, including positioning the mucosal side up and down, turning off the circulating fluid, as well as applying stretch. In conditions defined as “no stretch,” extreme care was taken to secure the colon to the dish with minimal to no stretch applied to the tissue, but because the colon is pinned in place, there are unavoidable changes in tension due to intrinsic movements of the tissue. Stretch was applied to colons in a subset of experiments in which the colon was lengthened in both directions (longitudinal and circular) by 150%. In experiments where the proximal and distal colon were separated, the colon was cut in half at the middle colon. In all cases, circulating fluid was slowly heated up to 34-37°C, and the colon equilibrated for 20 min prior to imaging.

### In situ Ca2+ imaging

Whole colons were removed from E2a-GCaMP6 mice, cut open longitudinally opposite to the mesenteric border, and pinned out in Sylgard-lined dishes as described above. Preparations were superfused with carboxygenated (95% O2, 5% CO2), warmed (34-37°C) ACSF and allowed to equilibrate for 20 min before imaging. GCaMP signals in myenteric neurons were recorded with an upright DM6FS Leica fluorescent microscope (Leica, BuffaloGrove, IL) and a Prime 95B Scientific Complementary Metal-Oxide-Semiconductor (CMOS) camera (Photometrics, RoperScientific, Tucson, AZ) using 10X or 20X objective lens, and images were collected with Metamorph software (MolecularDevices, San Jose, CA). To determine activity patterns that repeated during CMC cycles, we collected seven 3-min movies over the same field of view imaged at ∼6-Hz sampling rate and 150-ms exposure time. For increased temporal resolution, we also collected shorter movies imaged at a 40-Hz sampling rate and 25-ms exposure time. ICCs and myenteric neurons were identified based on morphology, anatomical location (within myenteric plexus or at the submucosa-circular muscle border), and the dynamics of their calcium transients. Tetrodotoxin (TTX) 0.5 micromol/L (Sigma-Aldrich, T8024) was dissolved in freshly oxygenated ACSF on the day of the experiment.

### Data Analysis and Statistics

Image files collected in Metamorph were exported to ImageJ (National Institutes of Health, Bethesda, MD). The amplitudes of GCaMP signals were analyzed and quantified as previously described^10, 19, 20^ by calculating dF/F0 as, where F is the peak fluorescence signal and F0 is the mean fluorescence signal at baseline. Neurons with transients with a signal amplitude >4SDs above baseline were considered “activity.” Tissue displacement was determined using a Template-Matching plugin in ImageJ, which quantifies movement along the x- and y-axes, representing the circular and longitudinal muscle, respectively. Statistical tests were performed in Excel (Microsoft, Redmond, WA) and GraphPad Prism (GraphPad Software, San Diego, CA). Data are expressed as mean ± standard error of the mean, where n represents number of mice used. Statistical tests are indicated in figure legends; significance was defined as P < .05.

